# DAR1: A Flexible Framework for Ecosystem Model Exploration

**DOI:** 10.1101/2023.11.23.568480

**Authors:** Barbara Duckworth, Stephanie Dutkiewicz, Oliver Jahn, Christopher N. Hill, Christopher L. Follett

## Abstract

Global scale biogeochemical ocean models, such as the MIT-Darwin model, are important tools to help us understand the mechanisms which drive elemental cycles and marine planktonic ecosystems. However, these models are difficult to use and their complex outputs can make them tricky to interpret. Here we present DAR1, an easy-to-use framework written in Julia that maintains all of the ecological and chemical complexity present in the MIT-Darwin model while removing the computational and physical complexity. We provide three studies to showcase DAR1 features: a framework for nutrient amendment bottle experiments; a test of steady state assumptions on plankton community structure at the global scale; and a test of theoretical predictions for the niche space of nitrogen-fixing plankton. DAR1 is ideal for model based testing of ecological predictions from field and experimental observations while also providing access to modeling tools for the non-expert.

## 1 Introduction

Marine microbes play an important role in biogeochemical cycling by fixing atmospheric carbon into organic compounds which, when exported to depth, lead to carbon sequestration in the deep ocean (Falkowski, 1994; Falkowski et al., 2003). The plankton community is equally important for feeding higher trophic levels and supporting the production of global fisheries. Phytoplankton community structure strongly influences these biogeochemical and ecological pathways, which are in turn affected by physical processes and nutrient availability (Dutkiewicz et al., 2020b; Caputi et al., 2019; Tréguer et al., 2018). Thus, understanding plankton community structure is best done in the context of the physical-chemical environment that they inhabit and in turn influence. Computational models can help us do this. The development of global ecological-biogeochemical ocean models has been critical for understanding the underlying relationships between the tightly coupled ecology and biogeochemistry of the marine environment (Dutkiewicz et al., 2009; Aumont et al., 2015; Ward et al., 2018). These “mechanistic” models are composed of differential equations describing the dynamics and interplay between the physics, biogeochemistry, and ecology of the ocean.

The MIT-Darwin model is one such model that was developed, in particular, to be able to flexibly model plankton distributions in the ocean and their interactions with global circulation and biogeochemical cycles (Follows et al., 2007; Dutkiewicz et al., 2015a). Past applications of this model have explored the drivers of plankton diversity (Dutkiewicz et al., 2020b); explained spatial and temporal patterns in species biogeography (Ward et al., 2014; Follett et al., 2018, 2022); and made predictions about how communities might be altered in a warming ocean (Dutkiewicz et al., 2019). MIT-Darwin follows an array of chemical species such as C, N, P, Fe, Si, and O_2_, that are described mathematically by a series of equations dictating their partitioning and transformation between living, detrital, and inorganic pools. For instance, a nutrient such as dissolved inorganic nitrogen will be taken up by autotrophic phytoplankton as a function of the local light and temperature environment. These phytoplankton, in turn, will be grazed by zooplankton and both will contribute dead organic matter to the detrital pool. MIT-Darwin contains all the equations and infrastructure to model a wide spectrum of plankton types and their interactions. The model takes a “trait based” approach. Plankton groups can be classified by a set of master parameters which are then scaled as a function of size. This approach has been used to simulate global communities from an handful of plankton (Dutkiewicz et al., 2015a) types up to hundreds (Dutkiewicz et al., 2020a).

MIT-Darwin is part of the MIT general circulation model (MITgcm, (Marshall et al., 1997)). The MITgcm physical configuration is highly flexible, allowing for different horizontal resolutions (from km to hundreds of km) in regional (e.g. (Bertin et al., 2023; Wilson et al., 2019)) and global (e.g. (Dutkiewicz et al., 2015a; Kuhn et al., 2019)) simulations, or as a 1-D water column (e.g. (Hickman et al., 2010; Wu et al., 2021)). Wind stresses, together with heat and freshwater fluxes set up mixing and circulation patterns (Forget et al., 2015; Marshall et al., 1997), and these flow fields advect and diffuse the chemical and biological “tracers” (e.g. the nutrients, plankton types, detrital matter etc) of MIT-Darwin.

This modeling framework, and others like it, has been extremely valuable as we seek to understand how marine ecosystems function and how they will evolve as the ocean warms. Unfortunately, all of this realism comes at a substantial cost in both computational resources, and the skills required to leverage them. Using the MIT-Darwin model requires niche knowledge and technical skills with a difficult learning curve. Like many global scale circulation and biogeochemistry models, it is written in Fortran, requires a complicated build environment, and any changes to the configuration or parameter choices involve changing numerous input files. Global scale simulations require large computer clusters and, depending on the spatial resolution of the configuration, can take hours to months to run for several years of model time. Complexities around both running the full MIT-Darwin model and interpreting the output of the simulations create a barrier to integrating observations with model predictions and testing new theoretical ideas. The number of scientists who can effectively use the model is small. One strategy to resolve these issues is to develop a framework that is consistent with the full MIT-Darwin model but is easier to operate and computationally inexpensive.

Here we present a new interface for a zero-dimensional configuration of the MIT-Darwin model, “DAR1” (see Figure 1), built using the Julia packages MITgcmTools and ClimateModels (Forget, 2024, 2022). DAR1 operates the MIT-Darwin model at full biochemical complexity, but within a single pixel/grid cell. These choices enable enhanced computational efficiency and a significantly simplified user experience. Changes in model parameters (see Table 1 for term definitions) and the specifics of the environment (e.g. temperature, nutrient, light) are set in a single easily edited file. Available documentation details the model setup, and a Docker container is provided to alleviate the build environment requirements (Merkel, 2014). Under many conditions, it takes only seconds to run on a standard desktop machine.

**Figure 1.**
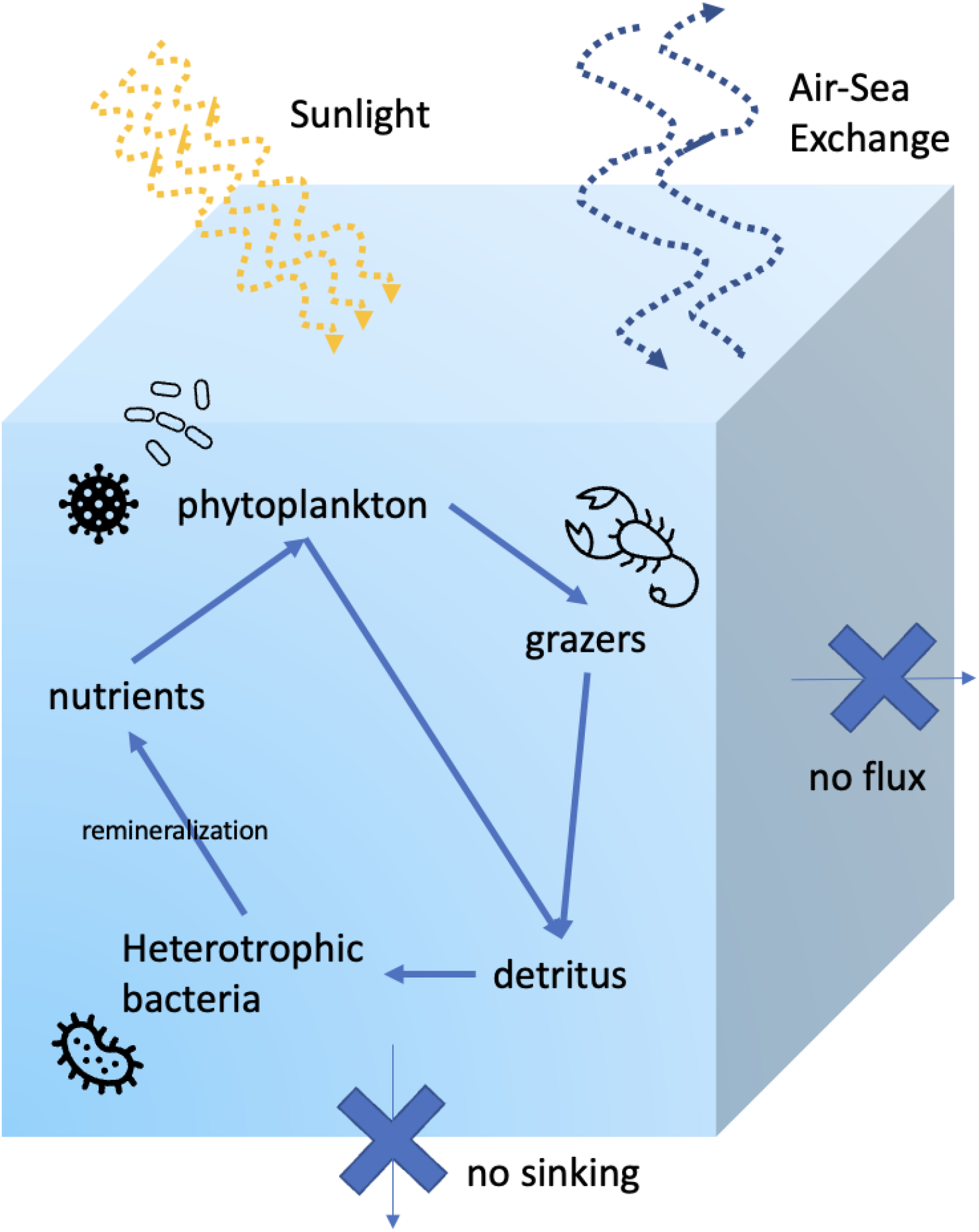
DAR1 Model Configuration: The DAR1 system models a single, well mixed parcel of water. The standard configuration consists of the MIT-Darwin ecosystem-biogeochemical model, with free gas exchange with the atmosphere. Horizontal and other vertical fluxes are set to zero. These conditions can be modified by the user.

**Table 1.** Definitions for technical terms used in the main text.

| Term | Definition |
| --- | --- |
| Tracer (State variable) | A term used in MIT-Darwin/DAR1 to describe all nutrients, biomass, and organic matter that change over time based on model calculations of losses and gains (synonymous with “state variable”). |
| Namelist | Fortran file type defining parameters and variables |
| Docker | A set of products that use OS-level virtualization to deliver software in packages called “containers”. |
| Docker container | A standardized, encapsulated environment that runs applications . |
| Docker image | A lightweight, standalone, executable package of software that includes everything needed to run an application: code, runtime, system tools, system libraries, and settings. It is used to build a container. |
| State variables (Tracers) | Fields representing biomass, nutrients, and organic matter whose values are updated at each timestep in the model based on calculated loss and gain terms (in MIT-Darwin this also includes advection and mixing). |
| Parameters | Values used in the model to define specific rates (e.g. maximum growth rate, grazing rates) or constants (e.g. maximum Chl to C) . |
| Build-time parameters | Parameters that must be set before the model is compiled, sometimes called “compile-time”. |
| Run-time parameters | Parameters with values that can be changed before or between simulations without recompiling the model. |
| Build | The compilation process, during which source code is converted to executable code. |
| Initial condition | The starting value of an evolving state variable, such as biomass or nutrient concentration. |
| Time step | The incremental change in time for which the governing equations of the system are being solved. |
| Steady state | All state variables have reached equilibrium, such that they no longer change from timestep to timestep. |
| NetCDF | A data format that supports labeled and annotated multi-dimensional scientific data. |

This paper describes the design and implementation of DAR1. It also provides three use cases: a nutrient amendment bottle experiment; a comparison of time-varying vs. steady state community predictions; and an exploration of the niche partitioning of nitrogen fixers. The final case study provides a novel scientific result. We are hopeful that DAR1 will aid in bridging the gap between observations, observationalists, models and modelers. Additionally, this class of model is well suited for large searches of model parameter space, and could aid in formally fitting ecological models to ocean transect data.

## 2 Design

### 2.1 Structure and Configuration

DAR1 is composed of three main parts:

1. A configuration of the MIT-Darwin model, written in Fortran, specifying a 1×1×1 meter cube of water for the simulation, along with default values for light, temperature, and biogeochemical tracers.
2. A Julia package for building, initializing, and running the MIT-Darwin configuration.
3. A Docker container that includes all necessary dependencies to run DAR1. Docker containers allow code that normally has specific hardware and system requirements to be run on any computer regardless of operating system. (See Table 1)

DAR1 is configured to be a single well-mixed box of water with air-sea exchange at the surface, no flux of matter through the sides, and no sinking. Within this box, in this specific configuration of the MIT-Darwin model (see Supporting Information), nutrients are cycled from inorganic to organic through photosynthesis and respiration, move up the trophic levels through grazing, and return to inorganic form by heterotrophic bacteria (Figure 1). This system is similar to a bottle experiment in which a quantity of water is collected from the ocean, closed off, and then community composition and nutrients are measured periodically as time passes. The model configuration can be modified to alter the number of plankton and to exclude grazers, bacteria, or other functional groups. While this paper presents a single configuration, alternative configurations are possible and will be made available in the future.

### 2.2 Workflow

DAR1tools is a package written in Julia that contains methods to build, run, and modify the MIT-Darwin model configuration. A standard workflow involves:

#### 2.2.1 Building

Traditionally, due to hardware and dependency differences, the MIT-Darwin model needs to be “built” (see table 1 for technical jargon such as this) on whichever computer will be running it. With DAR1, the build step is only done once, unless build-time parameters for the model need to be changed. The Docker container comes with the model already built and fit for the specifications of the docker container, allowing Docker users to avoid this step.

#### 2.2.2 Setup

Each experiment is set up with a unique identifier, called a run ID. This setup process creates a folder to store the simulation runs, links up the MIT-Darwin model code, copies over initial parameter files (see “namelist files” in Table 1), and returns an object that is used in all following steps to specify which files to modify and run.

#### 2.2.3 Parameter modification

Once the experiment is set up, the namelist files that contain run-time parameters can be modified. DAR1 has ready-made functions to change the most commonly accessed parameters, such as specifying the length of run, temperature, and initial nutrient and biomass concentrations. These ready-made functions cover the basics, but the user can also change any runtime parameter if the namelist file, group, and parameter names are known as per the MIT-Darwin documentation (Supporting Information). Parameters are often “seeded” from values taken from a global MIT-Darwin run. Therefore, DAR1tools contains a sample netCDF file with typical global tracer values for a one year time period at weekly temporal resolution.

#### 2.2.4 Running the model

After setup and parameter modification, the simulation can be run using a single command. The runtime for a simulation depends on how frequently output files are written. On a Lenovo Thinkpad L13 Core i5-1235U 8GB Memory, the three case studies from this manuscript took 45 seconds, 3 hours, and 45 minutes respectively.

#### 2.2.5 Examining results

The simulation output is located in the run ID folder, in the form of netCDF files (see Supporting Information). The output can be analyzed with Julia or another language. Example Julia and ipython notebooks with data visualizations are included in the DAR1 Github.

### 2.3 Parallelization

The MIT-Darwin model can be configured to run in any 3D arrangement of grid cells. Traditionally, tracers would be advected and diffused between grid cells to simulate the flow of currents in ocean basins. DAR1 takes advantage of MIT-Darwin’s ability to run many water “parcels” in parallel by turning off communication between grid cells, so each is its own isolated environment. This allows for a particular build of DAR1 to be run multiple times with varying initial conditions, such as increasing nutrient supply or seasonal temperature variations. This functionality is showcased in case studies 4.2 and 4.3 where a grid is used to explore global spatial variations in the former and a range of NO3 and PO4 initial conditions in the latter.

### 2.4 Constant Nutrients or other Parameters

In some cases, it is useful to set the concentrations of different tracers (e.g. nutrient concentrations) to be constant, rather than a time-changing quantity. In the default configuration of DAR1, all nutrients are cycled within the system – only gases can escape through the surface of the box. With DAR1, you can optionally choose to hold a number of nutrients or other model state variables (e.g. biomass, organic matter) constant. In the model, this is accomplished by asserting that the change between time steps is zero for the specified tracers.

## 3 Case Studies Demonstrate Functionality

DAR1 is built with a set of core functionalities which allow it to effectively bridge the gap between field observations and model predictions. In its simplest configuration, DAR1 operates like an ideal bottle experiment. The initial conditions are set from a parcel of water, and then the ecology and biogeochemistry inside the box are allowed to evolve, obeying the laws of mass conservation. A simple implementation of this use is explained in case study 4.1. Two additional functions of the model are explored in case studies 4.2 and 4.3. Case study 4.2 leverages the grid-based functionality of the MIT-Darwin model to compute the steady state across a grid of initial conditions taken from the surface of a modeled global ocean. Case study 4.3 uses this gridded functionality to search nutrient space for the niches of nitrogen fixers in the model ecosystem. Additionally, case study 4.3 introduces the ability of DAR1 to arbitrarily set the concentration of any tracer to a constant value over a timeseries. This allows the steady state to be explored in ecosystems that do not conserve nutrient concentration. In general, this functionality allows users to build generalized “chemostat” type models within the MIT-Darwin ecosystem.

## 4 Case Studies

### 4.1 A simplified, nutrient amendment, bottle incubation experiment

Here we show how DAR1 can be used to conduct a simulation of a bottle incubation experiment, using the North Pacific Ocean as a model environment. In a bottle experiment ambient seawater is collected from the environment and then manipulated (added nutrients, changing light, etc.) to understand the state of a planktonic community. For example, the oligotrophic gyre north of Hawaii is in a perpetual state of nitrogen limitation (Karl et al., 1997), with nitrate and deep seawater amendments leading to large increases in biomass and primary production in bottle experiments (Mahaffey et al., 2012). A similar experiment is run in DAR1 by taking surface water from the gyre just north of Hawaii, which has a NO_3_ concentration of 0.007 mmol N/m^3^, and increasing the nitrate 10x to a concentration of 0.07 mmol N/m^3^ (Figure 3). The “bottle” is then sampled every 12 hours over the course of a six day experiment. The biomass timeseries is then compared with the control experiment in which no additional nitrate was added. As expected in a nitrogen starved environment, we see that nitrogen amendment leads to a steeper increase in biomass over the first day, and an overall greater amount of biomass over the six day sampling period. The decrease in biomass after the peak is due to the re-partitioning of nutrients into dissolved and particulate organic matter.

**Figure 2.**
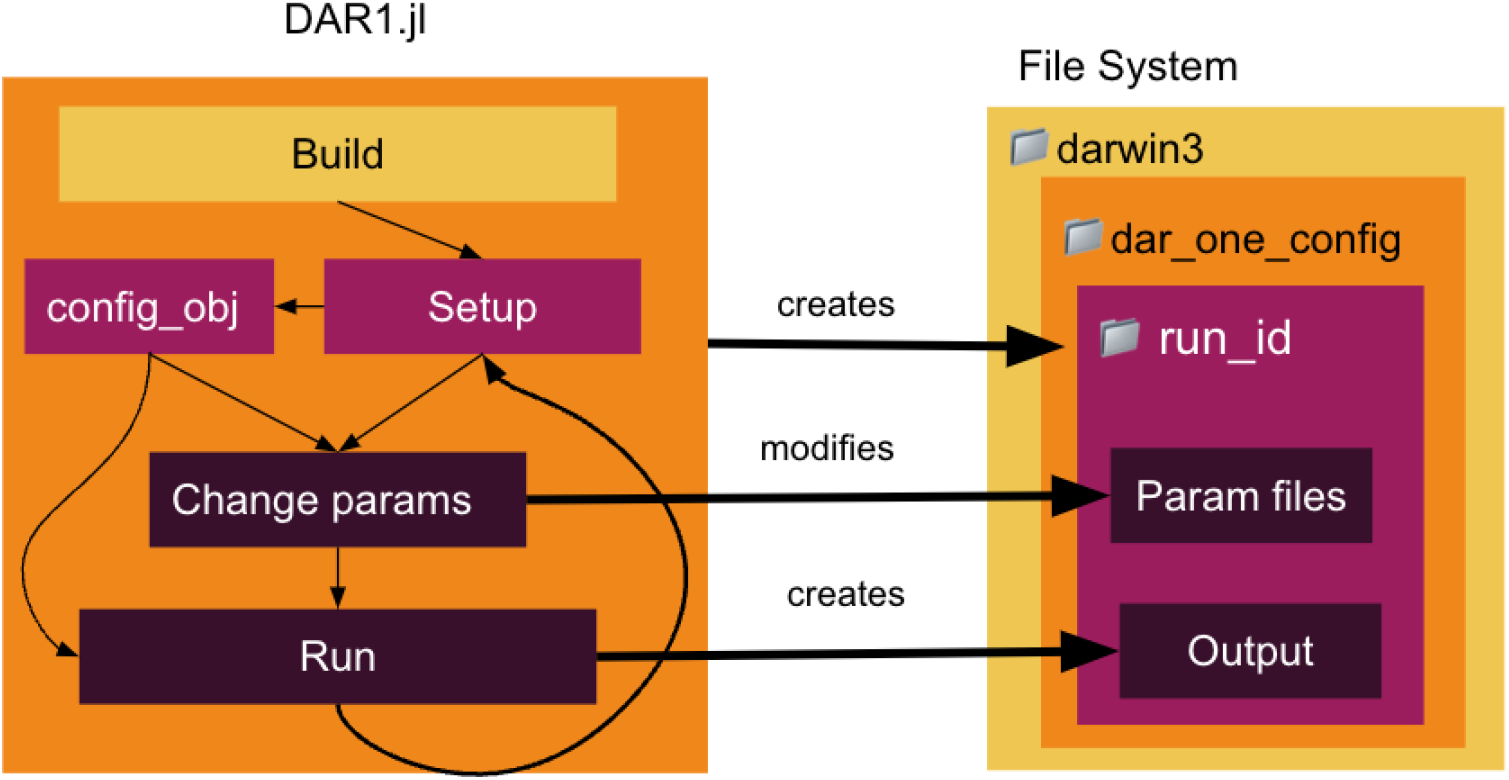
DAR1 Anatomy: DAR1tools provides an interface for interacting with the complex parameterization and file structure of the MIT-Darwin model.

**Figure 3.**
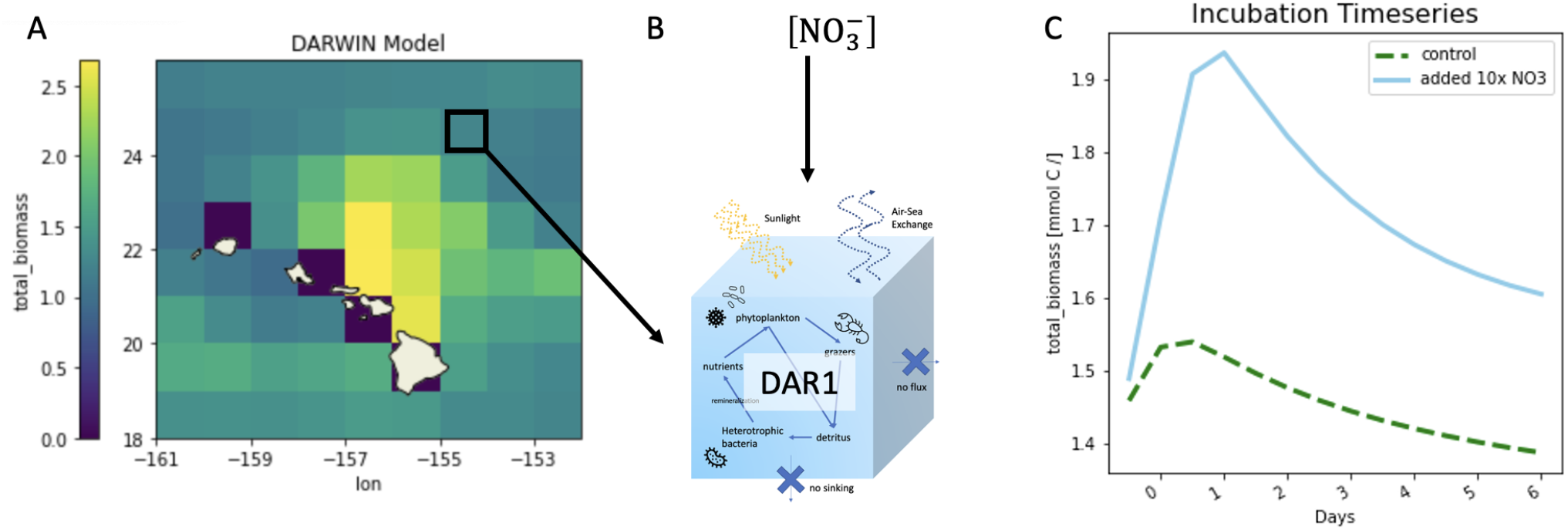
Case Study A: (A) Nutrients, phytoplankton, and other environmental variables were chosen from the point indicated on the map north of Hawaii to initialize DAR1. In the control, all variables were kept the same. The initial amount of NO_3_ was 0.007 mmol N/m^3^. In the nutrient amendment experiment, the amount of initial NO_3_ was increased to 0.07 mmol N/m^3^. (C) Shows the total amount of biomass in the control (green dashed line) and the case with additional nitrogen (solid blue line). In the case with additional nitrogen, the total biomass remains greater than that in the control.

This type of time series simulation is a standard work-flow with DAR1 and is a useful analogue to real world in-situ experiments. Users can pick an initial condition and use DAR1 to simulate the temporal evolution. A link to a step-by-step tutorial for replicating this case study is available in the Supporting Information.

### 4.2 Community Structure: Time Varying vs. Steady State

Ecological theories are powerful tools to explore the underlying mechanisms setting organism distributions and co-existence (Tilman and Sterner, 1984; May, 1972; Armstrong, 1999). With elegant and simple equations the fundamentals of the system can be explored, providing unique insight. However, there remains a question about when the simplicity is too much given the vastly more complex real world, especially when often these theories are solved assuming a steady state: a situation when the system does not change over time.

Previous MIT-Darwin model studies have utilized steady state theories (e.g. Resource Competition Theory, Resource Ratio Theory (Tilman, 1977; Tilman and Sterner, 1984)) to understand the controlling mechanisms on the resulting modeled plankton community structure (Dutkiewicz et al., 2009, 2020b; Ward et al., 2014). Here, we demonstrate that (as found in (Dutkiewicz et al., 2009)) steady state predictions are helpful for understanding oligotrophic gyres, but are less insightful at high latitudes. In particular, with DAR1 we can investigate what the actual steady state results would be without using the full 3D global configurations. In this setup we implement a parallel version of DAR1 (i.e. many simulations run at the same time) inside Docker.

Over time, species abundance will eventually converge to, or oscillate around, a steady state value. This property can be understood using a system of Monod equations (simplified from what is in DAR1) describing the total biomass of phytoplankton, *P*, and its limiting nutrient, *R*, supplied to the system at rate *S*_*R*_:

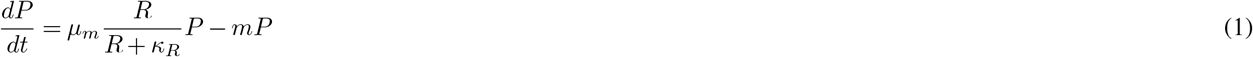

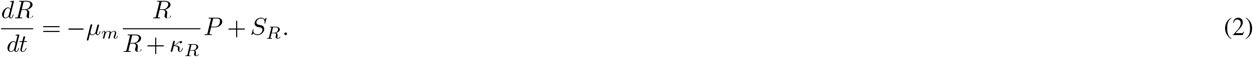

Here, *µ*_*m*_ is the maximal growth rate, *κ*_*R*_ is the half-saturation constant, and *m* is a loss term representing death, sinking, grazing, etc. The system reaches a stable equilibrium if the derivatives go to zero 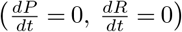. Fluctuating steady states can also be reached where the long-time average of the derivatives is zero. For the above simplified system of equations, a constant steady state is reached where the community composition is uniquely determined by the steady state nutrient concentration (Tilman and Sterner, 1984; Ward et al., 2014; Dutkiewicz et al., 2020b). However, this leaves the question of when the predictions of the steady state theory can be applied to the spatially and temporally fluctuating ocean. With DAR1, we can explicitly ask this question in the context of the full MIT-Darwin ecosystem by taking a vector of initial parameters (nutrient concentrations, biomass, light, temperature) and calculating the abundance of phytoplankton and nutrients over time using the same equations that power the MIT-Darwin model. With no fluctuations in light or temperature, and without transport from outside the parcel of water, the system can be run to steady state (i.e. when biomass, nutrients no longer change over times) or fluctuating steady states where the long-time average of the derivatives is zero.

For this case study, DAR1 is configured to run in a 160×360 matrix (each member of the matrix matching a 1 degree grid cell of a global configuration of MIT-Darwin), with no flux between cells. The nutrient and biomass concentration are initialized using the yearly average temperature, light, nutrient concentrations, and biomass of each cell’s counterpart in the global MIT-Darwin model. This matrix of DAR1 simulations are run for 10 years each, until the plankton communities reach a steady, or quasi-steady, state. We average over the last year of the simulation to get the “steady state” prediction from DAR1 and compare this output to the global fully dynamic MIT-Darwin simulation. We use the Bray-Curtis Dissimilarity Index defined as (Bray and Curtis, 1957):

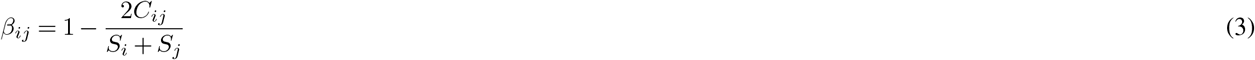

where

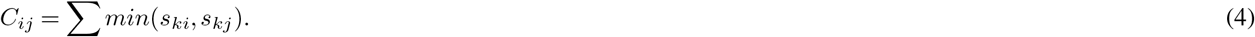

The index variables *i* and *j* refer to the two simulation setups being compared (the fully dynamic simulation and the matrix of DAR1 simulations) and the *k* index refers to the plankton types. Small *s*_*ki*_ is thus the biomass of a single type *k* from simulation *i* and *S*_*i*_ is the total biomass for simulation *i* (the same holds for index *j*). *C*_*ij*_ is the sum of the minimum biomass over all plankton types between the two simulations *i* and *j*. High dissimilarity indicates significant differences between the compared community structure. Conversely, a low dissimilarity score indicates that the plankton communities are similar. We compute the *β*_*ij*_ for each DAR1 steady state run against the corresponding geographic location of the MIT-Darwin dynamic model.

This comparison (Figure 4) indicates that oligotrophic gyres have a low dissimilarity score, meaning that the yearly average community of the global dynamic MIT-Darwin model is similar to the same conditions run to steady state over many years, as done in DAR1. In these regions, resource competition theory can be useful for understanding the structure of the phytoplankton community. In nutrient replete, high-latitude, waters the dissimilarity score is high, indicating that the community at steady state is not an appropriate approximation of the more realistic global model, which includes seasonal environmental changes and as such dynamically varying community structure. For these regions, the dominant driver of phytoplankton community composition is not simply an organism’s ability to draw down nutrients below competitor’s requirements as suggested by resource competition theory. Rather, it is the interplay between the time-varying forcing, biotic interactions, and the equilibration timescales of the ecosystem. A link to a tutorial for running this case study can be found in the Supporting Information.

**Figure 4.**
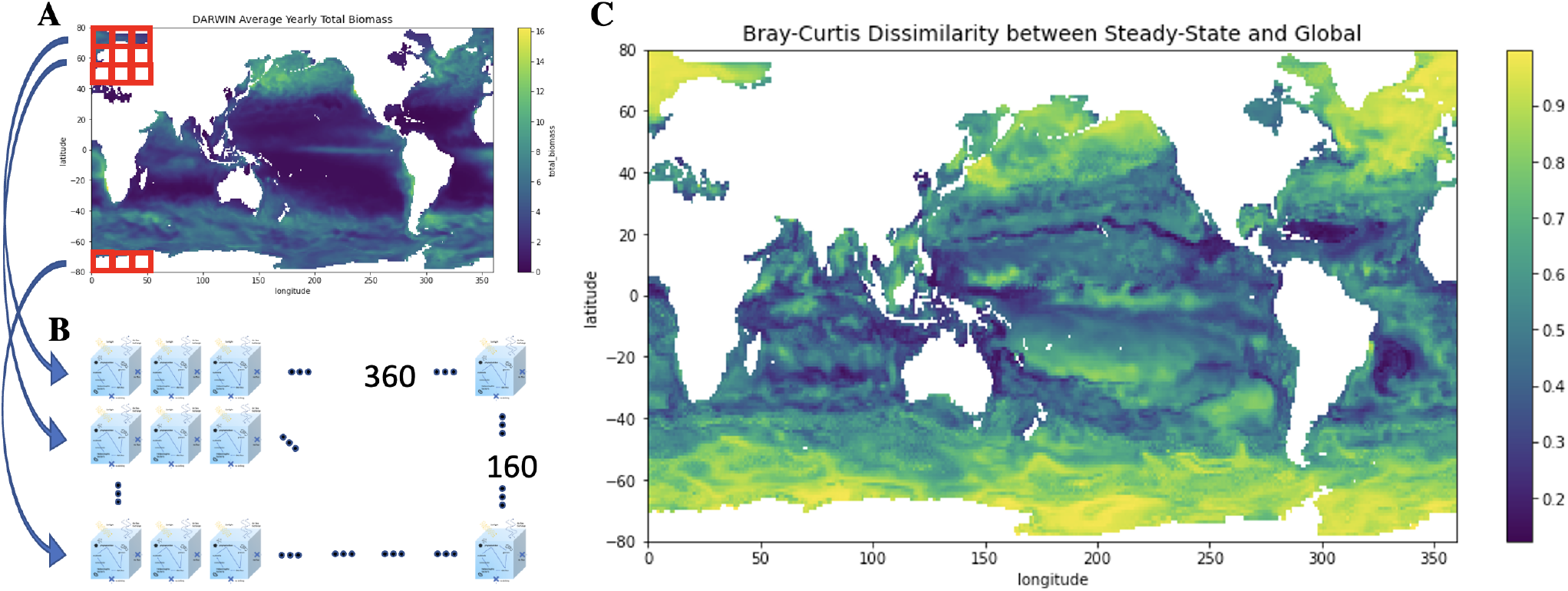
Overview of Case study B: (A) The average total biomass over the course of a year in the MIT-Darwin model. Variables (such as nutrients, plankton type abundance, sunlight, and temperature) from the dynamic, full 3 dimensional MIT-Darwin model are used to initialize a matrix of DAR1s, shown in (B). The DAR1 grid is 360×160 large, with each DAR1 “cell” corresponding to a lat/lon point in the dynamic 3-dimensional MIT-Darwin model. The grid of DAR1s is run 10 years forward, with final “steady state” values calculated from the average of the last two years. (C) The Bray-Curtis Dissimilarity index between the steady state community versus the annual community from the dynamic MIT-Darwin model simulation. The steady state is highly dissimilar to MIT-Darwin in regions with high values (yellow), and low values (blue) indicate areas where the community structure of the two setups are comparable.

### 4.3 Resource Ratio Theory: Niches of Nitrogen Fixation

Nitrogen fixers or diazotrophs can fix nitrogen gas into an organic form, rather than relying on the often low concentrations of fixed nitrogen such as nitrate (Zehr, 2011). This makes them unique and important members of the microbial community (Zehr and Capone, 2020). In general, these organisms have slower growth rates than similar sized non-diazotrophs given the energetic constraints of fixing nitrogen. Several studies have explored the environmental factors defining the biogeography of photosynthesising diazotrophic plankton (Monteiro et al., 2011; Dutkiewicz et al., 2014; Follett et al., 2018). Using a combination of resource supply ratio theory (Tilman, 1977) and numerical (MIT-Darwin) model simulations these studies suggest that diazotrophs co-exist with faster growing non-diazotrophs where the supply of iron and phosphate is in excess relative to the non-diazotrophs’ requirements (Ward et al., 2013; Dutkiewicz et al., 2012). Here, we use DAR1 to explore these ideas in more detail.

In this experiment, we run a series of simulations to steady state across a range of nitrate and phosphate concentrations (see Fig. 5). For simplicity, we set iron to be in excess, so it will never limit the growth of plankton in the system. We also fix DIN and PO_4_ concentrations; though this does violate mass conservation, this flexibility in DAR1 allows us to directly explore nutrient ratios that are difficult to do in dynamically evolving nutrient fields. We use concentrations of biomass and organic matter from north of Hawaii as initial conditions (Fig. 3). Temperature (24^*°*^ C) and light (average light at 50m, station ALOHA) are also held constant. More details on the setup can be found in the Supporting Information and Tutorials.

**Figure 5.**
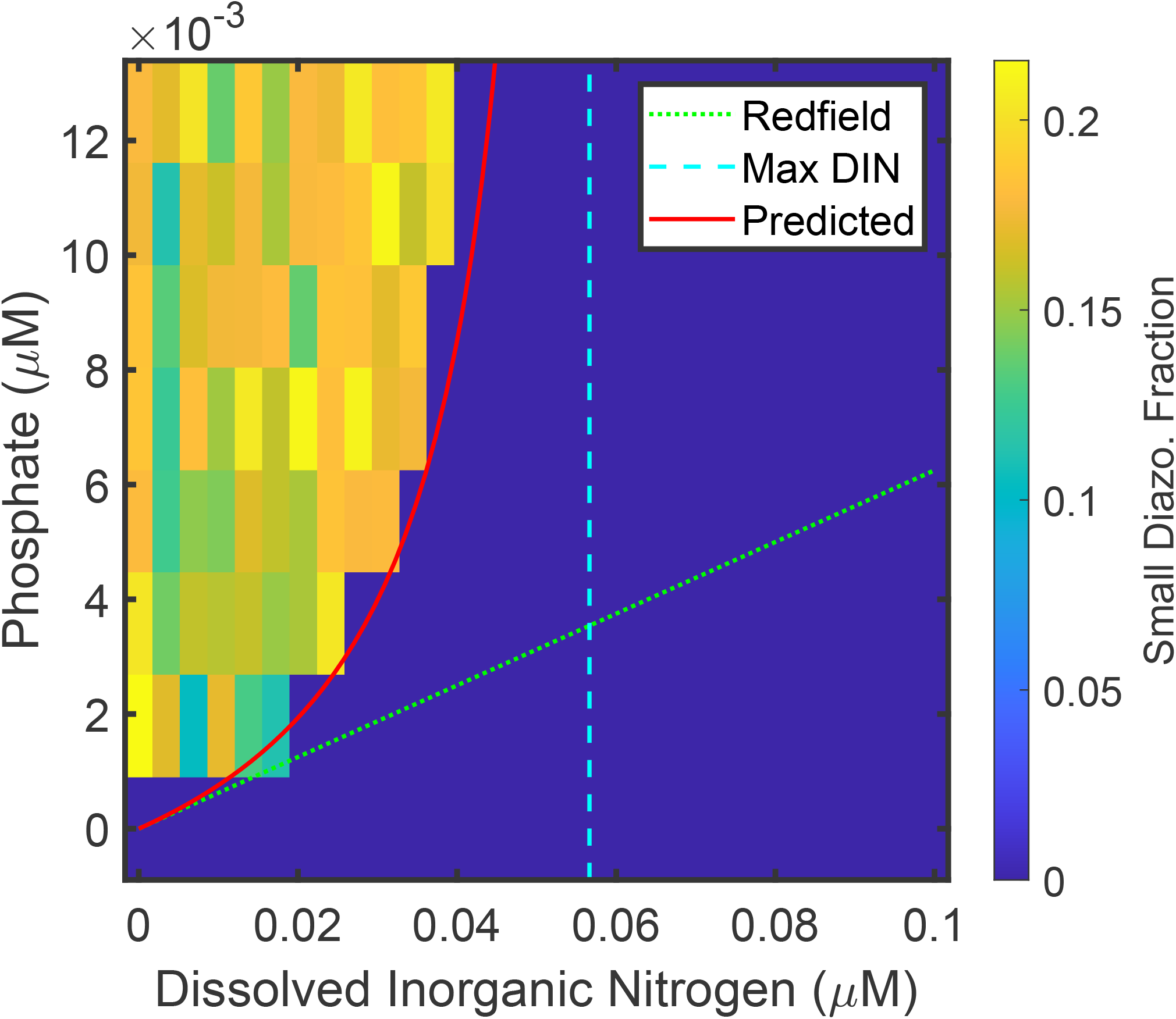
Overview of Case Study C: The steady state biomass fraction of the smallest nitrogen fixing diazotroph in MIT-Darwin is plotted as a function of phosphate and nitrate concentrations in the model. The solid red line is the predicted boundary where the small diazotrophs are expected to emerge (Eq. 6). The vertical dashed line is the maximum DIN value which allows for these diazotrophs (Eq. 7). The Redfieldian stoichiometry line, P:N=1:16, is also plotted (green dots). Increasing the amount of phosphate in the system leads to more diazotrophs whereas excess nitrate selects against nitrogen fixing organisms. The emergence of nitrogen fixers always occurs above the Redfieldian boundary. The results are shown here for the smallest of the diazotrophs for clarity, but similar arguments hold for all the size classes in the model.

Consistent with resource ratio theory (Tilman, 1977; Ward et al., 2013), we find the maximum diazotroph biomass when there is high phosphate and low nitrate conditions (Fig. 5). When the ratio of phosphate to nitrate is high, diazotrophs are able to co-exist with non-diazotrophs which are limited by nitrate concentrations. A natural hypothesis is that the exclusion/inclusion of diazotroph biomass would follow a boundary line with a slope of 1 : 16 given that this is the stoichometry we have set for the non-diazotroph. Instead, we find a more nuanced result (the curve pattern in Fig. 5b) suggesting that substantially more phosphate is required to allow nitrogen fixers to emerge as nitrate concentrations increase.

This result can be understood by thinking about competitive exclusion. In steady state, we expect that two organisms (limited by different nutrients) can only coexist if they have the same net growth rate. If mortality processes are similar, when a non-diazotroph in the model competes with a diazotroph of the same size the coexistence boundary should occur when the non-diazotroph’s nitrate limited growth rate is equal to the diazotroph’s phosphate limited growth rate. Mathematically, approximating growth as purely Monod, this will occur when

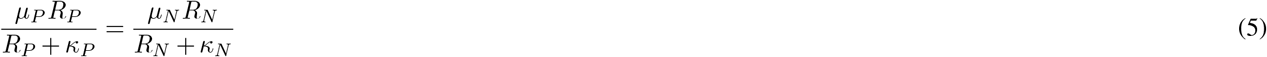

where the subscripts *P* and *N* refer to phosphate and nitrate respectively and the nitrate limited organism is the non-diazotroph.

Thus, *µ*_*N*_ and *µ*_*P*_ are the maximum growth rate for the non-diazotroph and the diazotroph. The half saturation constants are expressed as *κ*_*N*_ and *κ*_*P*_. In the model the growth rate of diazotrophs is less than the non-diazotrophs, i.e. *µ*_*N*_ *> µ*_*P*_. We use Eq. 5 to solve for the phosphate concentration that allows the coexistence of the two types:

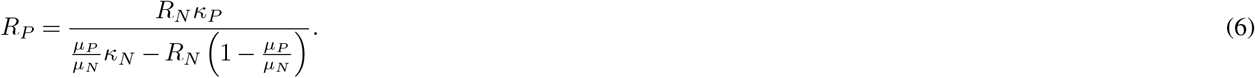

This solution suggests that as nitrate increases, the phosphate concentration required before diazotrophs can co-exist becomes larger and larger. At a certain concentration of nitrate, there is no phosphate level sufficient to sustain diazotrophs. This is expressed mathematically when the denominator in equation 6 becomes negative. This concentration is

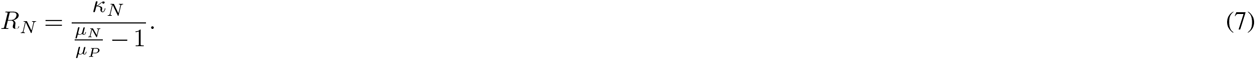

The slope of the boundary 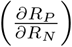 for low concentrations of *R*_*N*_ and *R*_*P*_ is

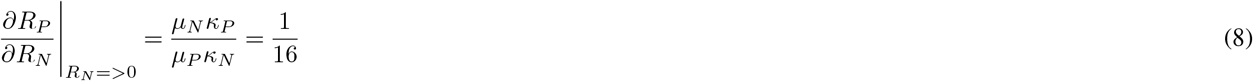

where the last step depends on the size scalings used in MIT-Darwin. Thus, the co-existence boundary is near the Redfield ratio at low nutrient concentrations before diverging for higher concentration. The predicted 1:16 line shown in Figure 5 forms the lower bound. The actual emergence curve is always above this line.

This case study demonstrates the usefulness of DAR1 in testing theoretical predictions. The simulations and calculations done here predict where different-sized diazotrophs should exist based on the observed nutrients, temperature, and light. This is a new result relative to previous work (e.g. Ward et al., 2013; Dutkiewicz et al., 2014). This insight, that is easy to explore in DAR1, provides a prediction that could be explored further in the full MIT-Darwin model and in laboratory experiments and the real ocean data.

## 5 Discussion

DAR1 bridges the gap between highly simplified proof-of-concept models and fully coupled ocean biogeochemical models by providing an intuitive interface for setting up and running a full complexity ecosystem simulation in a 0-D environment. DAR1 accomplishes this in two ways: computational efficiency and ease of use. DAR1 gains its efficiency by sacrificing physical complexity, but retains the full ecosystem of the more complex model. The ease of use comes from both an external interface that groups and simplifies the modification of large numbers of Fortran files, and by providing an easy way to modify model parameters and environmental conditions.

As with any tool, the effectiveness and impact of DAR1 hinges on whether the power it provides is correctly balanced by its ease of use. With DAR1 we have started with the full complexity and power of the fully coupled MIT-Darwin model (Dutkiewicz et al., 2020b; Marshall et al., 1997). From this we reduced the physical complexity of the model. With this sacrifice, we have provided a series of advantages which we hope will allow for broad use. Among the advantages are: platform flexibility; enhanced computational speed; and ease of operation. Often, it is innovations in ease of use which allow old ideas to be used effectively by practitioners in the field. Ocean Data View (Schlitzer, 2002), for example, allowed chemical oceanographers to optimally interpolate non-regular ocean transect data so that global patterns could be observed and compared effectively. The act of generating section plots from field data became a straightforward operation and allowed for field advances because of the ability to communicate patterns effectively. Modern ocean database projects seek to provide easy access to many datasets and the ability to co-localize and analyze them together (Ashkezari et al., 2021). DAR1 seeks to fill this role in the modeling world by connecting the output and conclusions of forward ecosystem models with observations and experiments.

The framework of DAR1, with its ease of use, provides a relatively easy introduction for non-experts into ecosystem modelling. But, we believe that simplified, yet fully integratable, models like this one will also be useful in testing the importance of ecological interactions in the ocean, formally fitting the ecological parameters used in plankton models, and aiding in the development of artificial intelligence techniques in the biogeochemical modeling sector. As shown with the case studies, the speed of use, ability to constrain the environment, and ecosystem structure allows for the exploration of theoretical constructs. We believe that DAR1 can be used for hypothesis testing, leading to ideas that can then be further explored in the full dynamic MIT-Darwin model and in the real ocean. Immediate plans for using this infrastructure include the testing of ecological drivers for seasonal range shifts in *Prochlorococcus* (Bian et al., 2023; Follett et al., 2022), exploring the importance of diel forcing on plankton populations (Tsakalakis et al., 2022), and exploring the importance of different grazing laws in setting community structure. Furthermore, we hope that structured output from simulations using DAR1 can help elucidate the situations under which statistical models for plankton populations are predictive (Bian et al., 2023; Bardon et al., 2021). The rapid technical development in marine ecosystem modeling has led to progressively more complex and realistic models whose power is coupled with increasing difficulty of use. We believe that models like DAR1 will help extend the power of these models for new applications and for new users.

### Code and data availability

MIT-Darwin code is part of the MITgcm ((Marshall et al., 1997), https://mitgcm.readthedocs.io/en/latest/). Code is available at https://github.com/darwinproject/darwin3, and documentation and equations at https://darwin3.readthedocs.io/en/latest/phys_pkgs/darwin.html. An introduction to DAR1 and links to the tutorials and case studies can be found at https://barbara42.github.io/Dar_One/build/getting_started/. DAR1 source code can be found at https://github.com/CBIOMES/DAR1. Code for the case studies can be found at https://github.com/CBIOMES/DAR1/tree/main/case_studies with required data. Please see short appendices for more information.

## Appendix A: Supporting Information

### A1 The Global Biophysical MIT-Darwin Ocean Model

MIT-Darwin code is part of the MITgcm ((Marshall et al., 1997), https://mitgcm.readthedocs.io/en/latest/). Code is available at https://github.com/darwinproject/darwin3, and documentation and equations at https://darwin3.readthedocs.io/en/latest/phys_pkgs/darwin.html.

The model resolves the cycling of C, N, P, Fe, Si, O2 through living, detrital and inorganic pools. It is developed to be flexible in the number and type of plankton that is resolves. It follows a “trait-based” approach as each plankton is assigned a number of traits that define its size, trophic status, resource requirements, thermal responses, and for phytoplankton, light affinity. The combination of traits set the plankton’s resource uptake and growth parameters. The model is highly flexible in the complexity of the ecosystem, where it can be configured to include 1 to 1000s of plankton. The plankton can be autotrophic, grazers, or remineralizers of organic matter, or some combination of trophic strategy (i.e. mixotrophic). The model can be set up for random assignment of traits (see e.g. (Follows et al., 2007; Dutkiewicz et al., 2009)), traits largely set by size alone (see e.g. (Ward et al., 2012)), traits focusing on functional type (see e.g. (Dutkiewicz et al., 2015b)), traits focused on trophic strategy (e.g. (Ward et al., 2012; Zakem et al., 2018)). The setup used in this manuscript follows (Follett et al., 2022) and includes 3 size classes of heterotrophic bacteria (spanning 0.4 to 0.9 *µ*m), 23 phytoplankton (spanning from 0.6 to 110 *µ*m, and including pico-cyanobacteria, pico-eukaryotes, calcifiers, nitrogen fixers, silicifers), 8 size classes of mixotrophs (from 6.6 to 140 *µ*m), and 16 size classes of zooplankton (spanning 4.5 to 1,636 *µ*m). Grazing is parameterized using a Holling II function (Holling 1972) and is size specific such that grazers can prey upon plankton 5 to 15 times smaller than themselves, with an optimal size of 10 times smaller (Hansen et al., 1997; Kiørboe, 2019).

### A2 Coding Environments

Docker uses OS-level virtualization to make applications work seamlessly across different environments through software packages called “containers”. Docker containers can be thought of as virtual machines, but they can share the OS kernel with other containers running as isolated processes, reducing the size of each individual container. A Docker “image” is the set of instructions needed to build and start up a container.

A Docker image is provided alongside the DAR1 code to make getting started as easy as possible. The Docker image, which will be used to start up a container in which DAR1 experiments run, is linked to on the github. The Docker container encapsulates the DAR1 code and all its dependencies. Notably this includes Julia and all the packages DAR1tools relies on, Fortran and its compilers, and all other libraries needed to run the MITgcm.

Using the Docker command line tools, the DAR1 image can be accessed with the following command: docker pull birdy1123/dar1. The most up to date image will be linked to in the top-level README of the Github. Further instructions on how to use Docker are included in the DAR1 “getting started” documentation (see links in next section).

### A3 Code Documentation, Access, and Case Study Tutorials

An introduction to DAR1 and links to the tutorials and case studies can be found in Github, at https://barbara42.github.io/Dar_One/build/getting_started/. DAR1 source code can be found at https://github.com/CBIOMES/DAR1. Installation and setup instructions are included in the documentation. Code for the case studies can be found at https://github.com/CBIOMES/DAR1/tree/main/case_studies, with step-by-step instructions on the documentation site. All seed files used in the case studies, generated from global MIT-Darwin runs, are included within the Docker.

## Author contributions

The project was conceived by CLF and SD. Project implementation and development was done by BD with coding support from CNH and OJ. Case studies and theory developed by CLF and SD. Writing done by BD, CLF and SD. Additional edits by CNH.

## Acknowledgements

We acknowledge funding from the Collaboration on Computational Biogeochemical Modeling of Marine Ecosystem (CBIOMES) Simons Foundation under grants: 549931, SD, OJ; 553242, C.L.F.; 827829, C.L.F.;

